# *C9orf72*- derived proline:arginine poly-dipeptides disturb cytoskeletal architecture

**DOI:** 10.1101/2020.10.14.338566

**Authors:** Tomo Shiota, Riko Nagata, Sotaro Kikuchi, Hitoki Nanaura, Masaya Matsubayashi, Mari Nakanishi, Shinko Kobashigawa, Kazuaki Nagayama, Kazuma Sugie, Yoshito Yamashiro, Eiichiro Mori

## Abstract

Amyotrophic lateral sclerosis (ALS) is an irreversible neurodegenerative disease caused by the degeneration of motor neurons, and cytoskeletal instability is considered to be involved in neurodegeneration. A hexanucleotide repeat expansion of the *C9orf72*, one of the most common causes of familial ALS, produces toxic proline:arginine (PR) poly-dipeptides. PR poly-dipeptides binds polymeric forms of low complexity sequences and intracellular puncta, thereby altering intermediate filaments (IFs). However, how PR poly-dipeptides affect the cytoskeleton, including IFs, microtubules and actin filaments, remains unknown. Here we performed a synthetic PR poly-dipeptide treatment on mammalian cells and investigated how it affects morphology of cytoskeleton and cell behaviors. We observed that PR poly-dipeptide treatment induce the degradation of vimentin bundles at perinucleus and dissociation of β-tubulin network. PR poly-dipeptides also lead to alteration of actin filaments toward to cell contours and strength cortical actin filaments via activation of ERM (ezrin/radixin/moesin) proteins. In addition, we found that PR poly-dipeptides promote phosphorylation of paxillin and recruitment of vinculin on focal adhesions, which lead to maturation of focal adhesions. Finally, we evaluated the effects of PR poly-dipeptides on mechanical property and stress response. Interestingly, treatment of PR poly-dipeptides increased the elasticity of the cell surface, leading to maladaptive response to cyclic stretch. These results suggest that PR poly-dipeptides cause mechanically sensitive structural reorganization and disrupt the cytoskeleton architecture.

## Introduction

Amyotrophic lateral sclerosis (ALS), a common neurodegenerative disease, causes muscle atrophy and weakness by progressive and selective loss of motor neurons [Rowland and Shneider, 2001]. A hexanucleotide repeat expansion in *C9orf72* is common in familial ALS with frontotemporal dementia [DeJesus-Hernandez et al., 2011], which induces autophagy [Webster et al., 2016] and defects nucleocytoplasmic transport [Zhang et al., 2018; Zhang et al., 2015] in ALS pathology. The *C9orf72* mutation produces toxic proline:arginine (PR) poly-dipeptides [Kwon et al., 2014]. PR poly-dipeptides penetrate cell membrane and localize to membrane-free organelles. PR poly-dipeptides also inhibit mRNA splicing and ribosomal RNA biogenesis, eventually causing cell death [Kwon et al., 2014], which target proteins with low-complexity (LC) domains, including cytoskeletal proteins [Lin et al., 2016].

The cytoskeleton is composed of three distinct species: actin filaments, microtubules, and intermediate filaments (IFs). Many neurodegenerative diseases show genetic abnormalities associated with cytoskeletal proteins. Neurofilament light-chain (NFL) encoded by the *NEFL* gene is the causative gene for Charcot-Marie-Tooth disease type 2E (CMT2E) and ALS [De Jonghe et al., 2001]. The accumulation of phosphorylated neurofilaments (NF) is a characteristic pathological finding of ALS [Leigh et al., 1989]. Cytoskeleton-related genes, such as *PFN1* and *TUBA4A* which respectively encode actin filaments and microtubules, are also identified as causative genes for ALS [Smith et al., 2014; Wu et al., 2012]. In addition, the morphology of actin filaments and microtubules is abnormal in these familial ALS mutations [Heo et al., 2018; Smith et al., 2014]. These data suggest that abnormalities of cytoskeleton are involved in ALS pathology. PR poly-dipeptides bind to vimentin and disassemble vimentin in vitro [Lin et al., 2016], and may induce morphological changes to the cytoskeleton.

The cytoskeleton, especially actin filaments, binds to focal adhesion proteins to maintain cell morphology and polarity and to modulate extracellular mechanics [Fletcher and Mullins, 2010; Hurtley, 1998]. Focal adhesion is the linkage between actin filaments and plasma membranes. It is a complex structure consisting of multiple proteins such as integrin, vinculin, paxillin, focal adhesion kinase (FAK), talin, zyxin, alpha-actinin, vasodilator-stimulated phosphoprotein (VASP), and other proteins [Abercrombie and Dunn, 1975]. Abnormal focal adhesions are also associated to neurodegenerative diseases such as Parkinson’s disease [Edwards et al., 2011] and Alzheimer’s disease [Leshchyns’ka and Sytnyk, 2016]. Vinculin is found in the Hirano body, a hallmark pathology of ALS [Galloway et al., 1987], which suggests that focal adhesion contributes to ALS.

Although these pathological findings in ALS suggest the involvement of the cytoskeleton and focal adhesion in the pathogenesis of ALS, the manner in which *C9orf72*-derived PR poly-dipeptides affect these structures remains elusive. In this study, we investigated that PR poly-dipeptides disrupted the architecture of the cytoskeleton and enhanced focal adhesion. We observed the morphological changes in the cytoskeleton and focal adhesion proteins by using fluorescens imaging. As these structures are involved in mechanical stress [Burridge and Guilluy, 2016], we evaluated the mechanical stress response by using atomic force microscopy (AFM) and cyclic stretch experiment.

## Methods

### Peptide synthesis

A synthetic peptide consisting of twenty repeats of the PR poly-dipeptide (PR_20_) with an HA tag at the carboxyl terminus was synthesized (SCRUM, Tokyo).

### Cell culture

Human osteosarcoma cells U2OS were cultured in Dulbecco’s modified Eagle medium (DMEM) high glucose with 10% fetal bovine serum (FBS) and 1% penicillin-streptomycin at 37▫ in 5% CO_2_. The U2OS cells were used for experiments after a one-hour treatment at 37▫ with a synthetic peptide consisting of PR_20_ (final concentration 10µM). Rat vascular SMCs (Lonza, R-ASM-580) were grown in DMEM with 20% FBS and 1x Antibiotic-Antimyotic (Thermo Fisher Scientific).

### Fluorescence imaging of cytoskeleton

The U2OS cells or rat vascular SMCs were fixed in 4% paraformaldehyde in Phosphate-Buffered Saline (PBS) at room temperature 15 min and permeabilized with 0.1% Triton X-100 in PBS for 10 min. The fixed cells were incubated with a blocking solution (5% bovine serum albumin in PBS with 0.1% Tween20) at room temperature for one hour. The cells were incubated with primary antibodies, vimentin (Santa Cruz Biotechnology, sc6260), vinculin (Abcam, ab129002), phospho-paxillin (Cell Signaling, 2541S), or phospho-ERM (Cell Signaling, 3726S) in the blocking solution at 4▫ overnight. Secondary antibodies (Thermo Fisher Scientific, A-21422, A-21429) were incubated at room temperature for one hour in the blocking solution. Alexa Fluor488-phalloidin (Thermo Fisher Scientific, A12379) and Alexa Fluor555-β-tubulin (Abcam, ab206627) were also incubated at room temperature for an hour in the blocking solution. Images were captured using the confocal microscope FV3000 (Olympus, Tokyo) or LSM 710 (ZEISS). Captured images were analyzed by ImageJ (NIH) and FIJI with QuimP plugin, which provided by University of Warwick [Baniukiewicz et al., 2018] and Tubeness plugin (htpps://imagej.net/Tubeness).

### Atomic force microscopy

Atomic force microscopy (AFM) measurements were performed using a NanoWizard IV AFM (JPK Instruments-AG, Germany) mounted on top of an inverted optical microscope (IX73, Olympus, Japan) equipped with a digital CMOS camera (Zyla, Andor) as described in a previous study by [Nagayama et al., 2019]. Prior to an AFM imaging of the surface topography and mechanical properties of U2OS cells in PR_20_-treated cells, the cells were adapted to a CO_2_-independent medium (Invitrogen) for 30 min at room temperature (25 °C). AFM quantitative imaging (QI) mode was used to obtain a force–displacement curve at each pixel of 128×128 pixels (100 µm ×100 µm of measured area) by a precisely controlled high-speed indentation test using rectangular-shaped silicon nitride cantilevers with a cone probe (BioLever-mini, BL-AC40TS-C2, Olympus, Japan). The test was performed at a spring constant of 0.08–0.10 N/m and a nominal tip radius of 10 nm. The QI mode measurements were performed within an h after the transfer of the specimen to the AFM. These high-speed indentations were performed until a preset force of 1 nN was reached. This typically corresponded to cell indentation depths of 300–400 nm. Cell elasticity was calculated from the obtained force–displacement curves by applying the Hertzian model (Hertz, 1881), which approximates the sample to be isotropic and linearly elastic. Young’s (elastic) modulus is extracted by fitting all force–displacement curves with the following Hertzian model approximation:

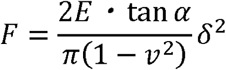

where F is the applied force, E is the elastic modulus, ν is the Poisson’s ratio (0.5 for a non-compressible biological sample), α is the opening angle of the cone of the cantilever tip, and δ is the indentation depth of the sample recorded in the force–displacement curves. Using the results of the Hertzian model approximation, we identified the Z contact points (specimen surface) and the elastic modulus of the specimens at each pixel and produced a surface topography map and elastic modulus map of the specimens.

### Cyclic stretch experiment

Cyclic stretch was performed using a uniaxial cell stretch system (Central Workshop Tsukuba University) as described in an existing study [Yamashiro et al., 2020]. The rat vascular SMCs were plated on silicon elastomer bottomed culture plates (SC4Ha, Menicon Life Science) coated with cell attachment factor containing gelatin (Thermo Fisher Scientific, S006100) and subjected to cyclic stretch with a frequency of 1.0 Hz (60 cycles/min) and 20% strain for six hours.

### Statistical Analysis

All experiments are presented as means ± SD. Statistical analysis was performed using Prism 8 (Graph Pad). The Mann-Whitney U test, a nonparametric test, was conducted. P < 0.05 denotes statistical significance.

### Data availability

All data supporting the findings of this study are available from the corresponding authors on reasonable request.

## Results

### PR_20_-treatment modulates the cytoskeleton architecture on U2OS cells

To investigate how PR poly-dipeptides affect the cytoskeleton, we first examined the morphological changes of vimentin in U2OS cells after exposure to 1μM of PR_20_ for an hour. Vimentin was predominantly localized at the perinuclear space, and bundles of vimentin were clearly detected in the control cells (CTRL; vehicle treatment) (Fig. 1A). Whereas in PR_20_-treated cells, the intensity of vimentin at the perinuclear space decreased, and the bundles of vimentin altered a mostly diffused background (Fig. 1A). The fluorescence intensity of vimentin at the perinuclear region was significantly reduced in PR_20_-treated cells (10.895 ± 5.192 a.u., n=57) compared to CTRL (14.616 ± 6.89 a.u., n=52) (Fig. 1B). Since vimentin affects microtubule polymerization [Shabbir et al., 2014], we next investigated the effect of PR poly-dipeptides on the morphology of β-tubulin. β-tubulin showed a filamentous network in cytoplasm and the cell peripheral region (Fig. 1C). On the other hand, it appeared as dots with most of the network structure diffused in PR_20_-treated cells (Fig. 1C). We evaluated the tube-like structures and quantified polymerized-microtubules using ImageJ with Tubeness plugin and measured the ratio of polymerized-microtubules on each condition. Compared to CTRL (0.15 ± 0.03, n=51), PR_20_-treatment significantly reduced polymerized-microtubules (0.11 ± 0.08, n=57; Fig. 1D). These results imply that PR poly-dipeptides induce the degradation of vimentin bundles at the perinucleus and dissociation of microtubule network.

**Figure 1.**
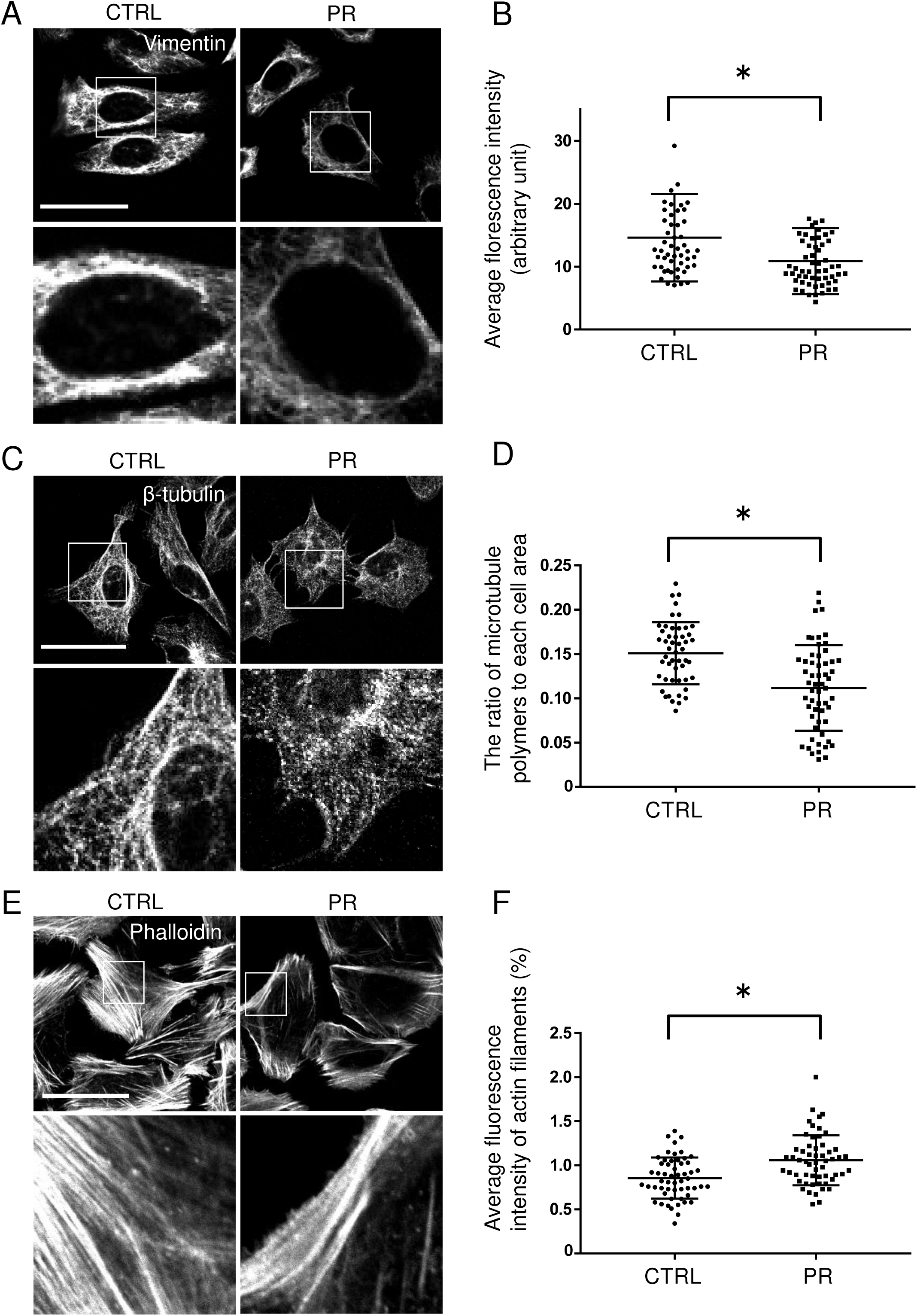
PR_20_ changes the architecture of cytoskeleton. Immunostaining of vimentin (A), β-tubulin (C) and Phalloidin (E) after treatment of 10 µM PR_20_ or untreated control. Scale bars are 50 µm. (B) Quantification of the fluorescence intensity of vimentin at perinuclear region. (D) The ratio of microtubule polymers to each cell area. (F) Average fluorescence intensity of actin filament in CTRL (n=55) and PR_20_-treated cells (n=56). *P<0.05, Mann–Whitney U test.

As vimentin and tubulin form a structural network with actin filaments [Jiu et al., 2015] [Morris and Hollenbeck, 1995], we further investigated the effect of PR poly-dipeptides on the organization of actin filaments. In the CTRL, actin filaments were extended straight across the whole cell-body (Fig. 1E) whereas after PR_20_-treatment, actin filaments disappeared from central and accumulated at the peripheral of cell-body (Fig. 1E). Strength of the actin filaments was evaluated by FIJI with QuimP plugin and measured its fluorescence intensity. PR_20_-treated cells showed high intensity (1.06 ± 0.28, n=56), compared to that of the CTRL (0.86 ± 0.23, n=55; Fig. 1F). These results suggest that PR poly-dipeptides lead to alteration of actin filaments toward to cell contours and strength cortical actin filaments.

### PR poly-dipeptides change in organization of actin filament and focal adhesion

To examine the formation of cortical actin filaments, we evaluated Ezrin/Radixin/Moesin (ERM) proteins, which cross-link the actin filaments with the cell membrane. Activated ERM proteins by phosphorylation are necessary for forming cortical actin and filopodia by polymerized actin filaments [Furutani et al., 2007]. In PR_20_-treated cells, ERM proteins were dramatically phosphorylated and upregulated compared to CTRL (Fig. 2A). Phospho-ERM proteins in PR_20_-treated cells were abundantly expressed and mainly localized to protrusive structures, such as filopodia, and not observed in the cytoplasm, while phospho-ERM localized only at the tip of the protrusion in CTRL cells (Fig. 2A). These data strongly support our findings (as presented in Fig.1E-F) that PR poly-dipeptides lead to the reorganization of actin filaments through an activation of ERM proteins.

**Figure 2.**
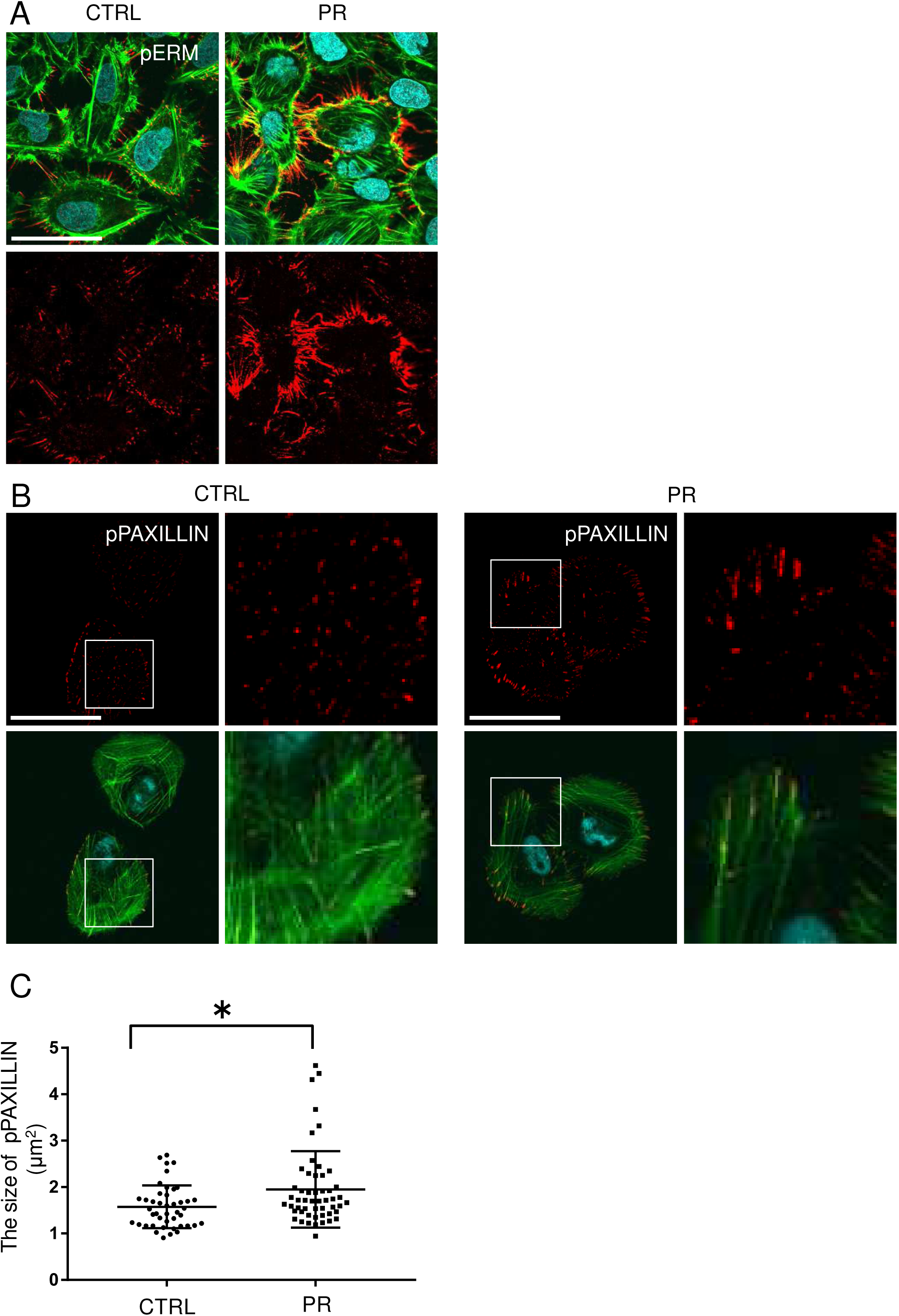
PR_20_ affects cytoskeletal actin binding proteins and focal adhesion (FA). A. Immunostaining of U2OS cells treated by 10 µM PR_20_ with phospho-ERM (red in A). B. Immunostaining of U2OS cells treated by 10 µM PR_20_ with p PAXILLIN (red in B), Phalloidin (green) and DAPI (blue) are shown. Scale bars are 50 µm. Focal adhesion size was evaluated by pPAXILLIN (in B) and quantified using ImageJ in CTRL (n=44) and PR_20_-treated cells (n=52) (C). *P<0.05, Mann–Whitney U test.

Alteration of actin filaments changes cell mechanics, thereby regulating cellular behaviour such as proliferation, migration and maturation of focal adhesions [Oakes et al., 2012; Parsons et al., 2010]. Phosphorylation of paxillin recruits to vinculin on the tip of the actin stress fiber, thereby making a hub between integrins and actin filaments and resulting in the maturation of focal adhesions. Mature focal adhesions form molecular complexes and grow in size [Gardel et al., 2008].

To investigate the effects of PR poly-dipeptides on maturation of focal adhesions, we evaluated the localization of vinculin, phosphorylation levels of paxillin and measured size of focal adhesions. Vinculin was predominantly expressed in nascent adhesions under nucleus in CTRL cells. However, it was localized at the cell periphery, which forms mature focal adhesions (Fig. S2A). In addition, phosphorylation levels of paxillin (pPAXILLIN) were detected at the periphery of PR_20_-treated cells, compared to CTRL cells (Fig. 2B). Size of pPAXILLIN was significantly larger than that of CTRL (Fig. 2C), even if the size of vinculin was comparable between the CTRL and PR_20_-treated cells (Fig. S2B). These observations imply that PR poly-dipeptides promote the phosphorylation of paxillin and recruitment of vinculin on focal adhesions, which lead to a maturation of focal adhesions.

### PR poly-dipeptides increase the elasticity of the cell surface

To further evaluate the changes of actin filaments and focal adhesions, we measured the elasticity of the cell surface by AFM. There were no significant differences in cell height between the two groups (3.87 ± 1.01μm, n=69 in CTRL, 3.993 ± 0.873μm, n=45 in PR_20_-treatment; Fig. 3A, B). Interestingly, the elasticity at cell center was clearly increased in PR_20_-treated cells (12.857 ± 8.196 kPa, n=45), compared to that of CTRL (8.562 ± 5.051 kPa, n=69; Fig. 3C, D). These results indicate that PR poly-dipeptides abnormally increase in cell stiffness.

**Figure 3.**
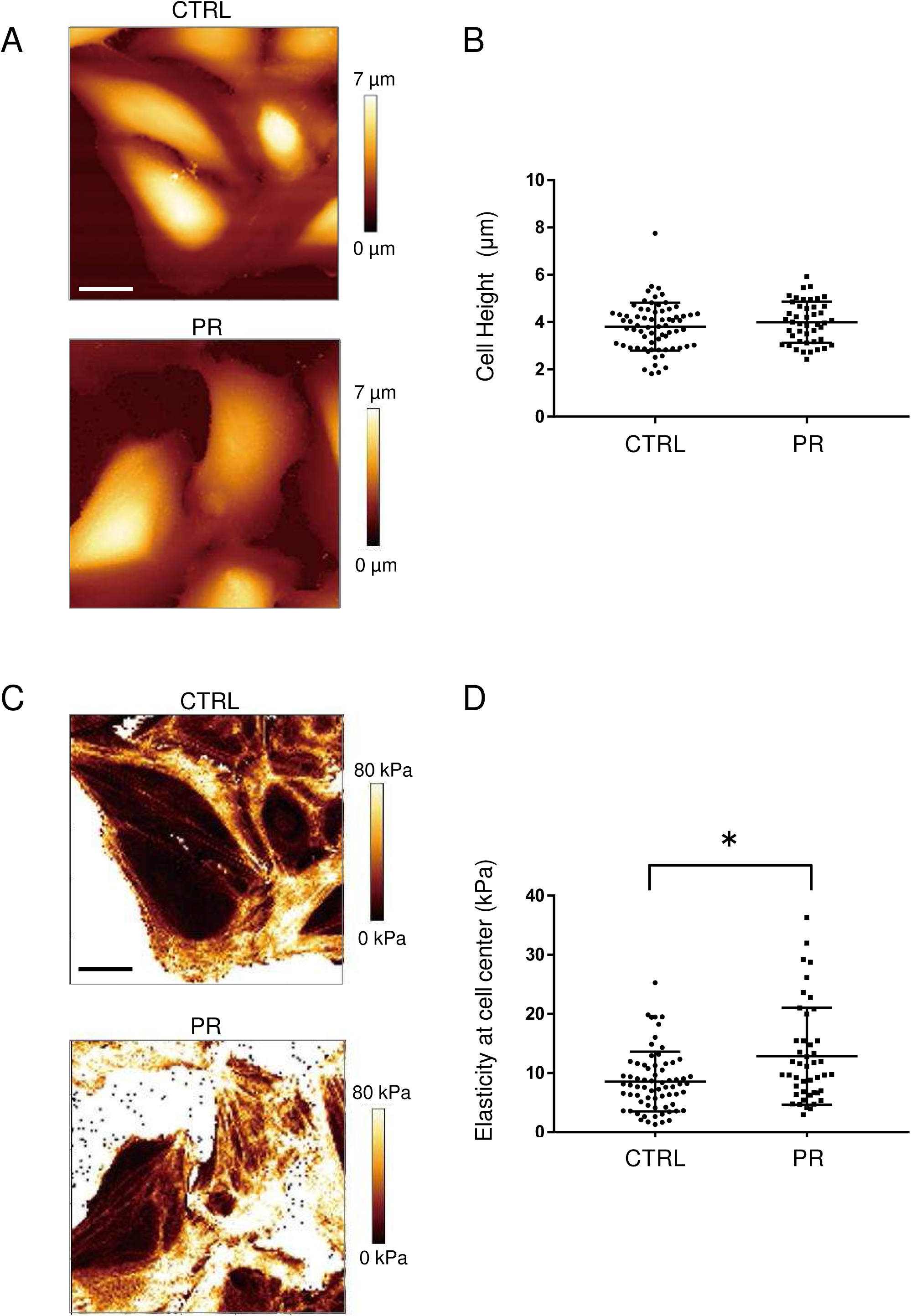
AFM images of the surface of U2OS cells. Representative surface topographic images (A) and elastic modulus maps (C) of U2OS cells measured by AFM. Scale bars are 20 µm. Quantification of cell height (B) and elastic modulus at cell center (D) in CTRL (n=69) and PR_20_-treated (n=45) are shown. *P<0.05, Mann–Whitney U test.

### PR poly-dipeptides attenuated the cyclic-stretch-induced reorientation of actin stress fibers

So far, we observed that PR-treatment induced the assemble of cortical actin and enhanced maturation of focal adhesions marked with increasing the size of pPaxillin and phosphorylation of ERM proteins. Mechanical force is transmitted via integrin and propagates into focal adhesion molecules to nucleus through actin filaments. Mechanical stress response is fundamental for maintaining cellular homeostasis, integrity and adaptation for pathological conditions. Therefore, we hypothesized that the PR poly-dipeptides-induced reconstitution of actin stress fibers alters mechanical stress response. To test this hypothesis, we examined the effect of PR_20_ on cyclic stretch-induced reorientation of actin stress fibers. Instead of U2OS cells, we employed rat vascular smooth muscle cells (SMCs) which are frequently used for cyclic stretch experiments. Rat vascular SMCs with or without PR_20_ treatment were subjected to cyclic stretch (20% strain, 1 Hz) for six hours. We then evaluated the orientation of actin stress fibers to the direction of cyclic stretch. As we expected, CTRL cells responded normally with a reorientation of actin stress fibers aligned to the perpendicular position, whereas PR_20_-treated cells failed to align correctly and decreased cell density after stretch (Fig. 4A). The histograms of the percentage of the orientation angle (θ) for each cell show that PR_20_ significantly suppressed the cyclic stretch-induced reorientation of stress fibers (17.0 ± 10.1°, n=96 in CTRL, 34.286 ± 23.104°, n=70 in PR_20_-treatment; Fig 4B). These results provide strong evidence that PR poly-dipeptides cause abnormal remodeling of mechanical stress response and disorganization of cellular homeostasis.

**Figure 4.**
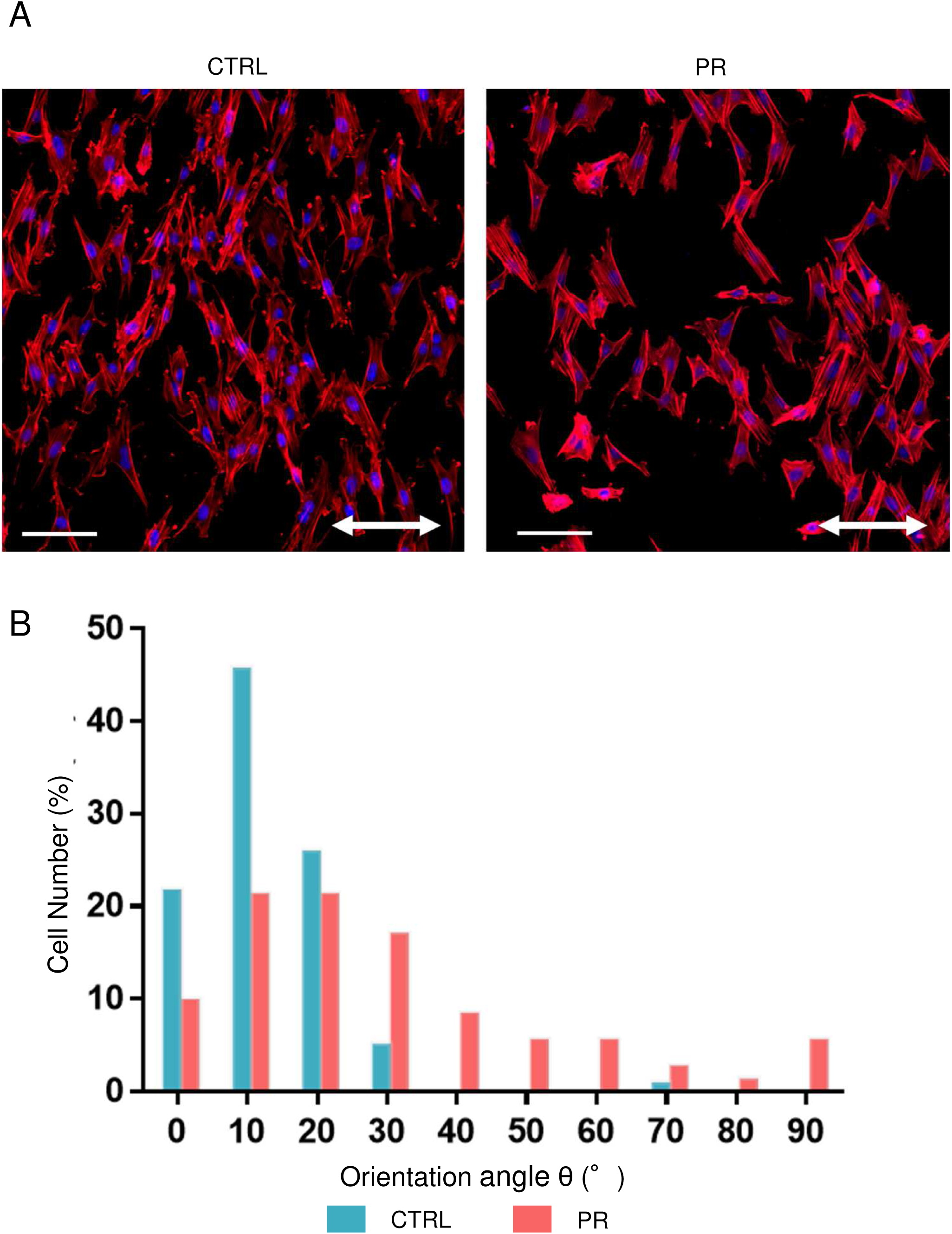
PR_20_ prohibits the cyclic-stretch-induced reorientation of actin stress fiber in rat vascular SMCs. Rat vascular SMCs with or without 10 µM of PR_20_ were subjected to cyclic stretch (20% strain, 1.0 Hz (60 cycles/min) for six hours. A. Two-way arrows indicate stretch direction. Phalloidin (red) and DAPI (blue) also shown. Scale bars are 100 µm. B. The orientation of each cell (bottom) was analyzed by measuring the orientation angle (θ) of the long axis of the ellipse relative to the stretch axis in CTRL (n=96) and PR_20_-treated cells (n=70).

## Discussion

The proper regulation of cytoskeleton architecture is important for many developmental and physiological processes in multicellular organisms [Hurtley, 1998]. Cytoskeletal proteins form a complicated network and reorganize in response to mechanical forces [Fletcher and Mullins, 2010]. In this study, we demonstrated how PR poly-dipeptides alter cytoskeletal morphology and induce maturation of focal adhesions and maladaptive response to mechanical stress.

In ALS, an inhibition of vimentin expression during neuronal development can cause motor neuron degeneration [Gomes et al., 2019]. Vimentin is essential for the early stage of neuronal development [Yabe et al., 2003] and initially forms the neuronal network [Giasson and Mushynski, 1997]. The degradation of vimentin by PR poly-dipeptides may prevent the acquisition of normal neuronal network, leading to neurodegeneration. PR poly-dipeptides also depolymerize microtubules (Fig. 1C). This result is consistent with microtubules depolymerization in the other familial ALS, *RAPGEF2* and *TUBA4A* [Smith et al., 2014; Heo et al., 2018]. Axonal transport, an important role of microtubules in neurons, is disrupted in *TUBA4A* mutation which contributes to dying back axonal damage and energy deficits in distal axons in ALS [Ferraiuolo et al., 2011], suggesting that PR poly-dipeptides treatment may also disturb axonal transport.

Actin filaments decrease in *PFN1* mutation [Sivadasan et al., 2016; Wu et al., 2012], and filopodia is increased in motor neurons of *SOD1* ALS mouse models [Osking et al., 2019]. As cortical actin induces neuronal differentiation [Flynn et al., 2012], the remodeling of cortical actin suggests that PR poly-dipeptides may inhibit axonal outgrowth and the normal differentiation of neurons. Filopodia is also required for synaptic connections after denervation [Osking et al., 2019]. PR poly-dipeptides might affect new synaptic connections via ERM activation.

We also showed that PR poly-dipeptides lead to abnormal maturation of focal adhesions (Fig. 2A). In a *SOD1* mutant mouse model, focal adhesions are strengthened and activated the astrocyte-associated pathway, leading to neurodegeneration [Lagos-Cabré et al., 2017]. PR poly-dipeptides also might induce neurodegeneration via signaling pathway mediated focal adhesion.

Further, our results show that PR poly-dipeptides change the mechanical stress response. When cytoskeletal rearrangement and redistribution of focal adhesions occur correctly, cell reorientation is observed along stretching direction [Ikawa and Sugimura, 2018]. The abnormal rearrangements of actin filaments and focal adhesions contribute to the suppression of cell reorientation. Increasing in cell stiffness which is regulated by actin organization and focal adhesions [Gauthier et al., 2012] is supported by maturation of focal adhesions in PR poly-dipeptides treatment. The distribution of actin filaments is also important for cell stiffness [Fletcher and Mullins, 2010; Smith et al., 2014]. Actin cortex forms a thin network with actin filaments and myosin motors within cell membranes. It regulates mechanical stress in the absence of stress fibers, and produces tension isotropically [Efremov et al., 2019], suggesting that PR poly-dipeptides increase cellular tension by cortical actin. Cell contraction by cytoskeleton causes neurite retraction in neurodegeneration diseases [Luo, 2002]. The stiffness of brain tissues also occurs with aging [Sack et al., 2011]. Cell stiffness may be an important key to loss of neuronal function.

In summary, this study observed that PR poly-dipeptides cause mechanically sensitive structural reorganization and disrupt cell homeostasis. PR-induced cytotoxicity is characterized by mechanical changes. The alternations of mechanical properties might be associated with in neurodegenerative diseases.

## Acknowledgments

The authors thank Keren-Happuch E Fan Fen for their critical reading of the manuscript.

## Figure legend

**Supplemental figure 1. Quantification of F-actin fluorescence intensity**. Schema of QuimP analysis procedure. Measurement of the fluorescence intensity of phalloidin using ImageJ. Automatically fit contours along the ROI drawn around the edge of the cell and measure the fluorescence intensity in the strip-shaped region of the width 0.7 µm inside the drawn contour.

**Supplemental figure 2. PR**_**20**_ **affects vinculin** (A) Immunostaining of U2OS cells treated by 10 µM PR_20_. Phalloidin (green), Vinculin (red) and DAPI (blue) are shown. Scale bar is 50 µm. (B) Focal adhesion size was evaluated by size of vinculin (in A) and quantified using ImageJ in CTRL (n=53) and PR_20_-treated cells (n=60). *P<0.05, Mann–Whitney U test.

## References

Abercrombie M, Dunn GA. 1975. Adhesions of fibroblasts to substratum during contact inhibition observed by interference reflection microscopy. Exp Cell Res 92:57–62.

Baniukiewicz P, Collier S, Bretschneider T. 2018. QuimP: analyzing transmembrane signalling in highly deformable cells. Bioinformatics 34:2695–2697.

Burridge K, Guilluy C. 2016. Focal adhesions, stress fibers and mechanical tension. Exp Cell Res 343:14–20.

De Jonghe P, Mersivanova I, Nelis E, Del Favero J, Martin JJ, Van Broeckhoven C, Evgrafov O, Timmerman V. 2001. Further evidence that neurofilament light chain gene mutations can cause Charcot-Marie-Tooth disease type 2E. Ann Neurol 49:245–9.

DeJesus-Hernandez M, Mackenzie IR, Boeve BF, Boxer AL, Baker M, Rutherford NJ, Nicholson AM, Finch NA, Flynn H, Adamson J, Kouri N, Wojtas A, Sengdy P, Hsiung GY, Karydas A, Seeley WW, Josephs KA, Coppola G, Geschwind DH, Wszolek ZK, Feldman H, Knopman DS, Petersen RC, Miller BL, Dickson DW, Boylan KB, Graff-Radford NR, Rademakers R. 2011. Expanded GGGGCC hexanucleotide repeat in noncoding region of C9ORF72 causes chromosome 9p-linked FTD and ALS. Neuron 72:245–56.

Edwards YJ, Beecham GW, Scott WK, Khuri S, Bademci G, Tekin D, Martin ER, Jiang Z, Mash DC, ffrench-Mullen J, Pericak-Vance MA, Tsinoremas N, Vance JM. 2011. Identifying consensus disease pathways in Parkinson’s disease using an integrative systems biology approach. PLoS One 6:e16917.

Efremov YM, Velay-Lizancos M, Weaver CJ, Athamneh AI, Zavattieri PD, Suter DM, Raman A. 2019. Anisotropy vs isotropy in living cell indentation with AFM. Sci Rep 9:5757.

Ferraiuolo L, Kirby J, Grierson AJ, Sendtner M, Shaw PJ. 2011. Molecular pathways of motor neuron injury in amyotrophic lateral sclerosiseditor^editors. Nat Rev Neurol. England, p 616–30.

Fletcher DA, Mullins RD. 2010. Cell mechanics and the cytoskeleton. Nature 463:485–92.

Flynn KC, Hellal F, Neukirchen D, Jacob S, Tahirovic S, Dupraz S, Stern S, Garvalov BK, Gurniak C, Shaw AE, Meyn L, Wedlich-Söldner R, Bamburg JR, Small JV, Witke W, Bradke F. 2012. ADF/cofilin-mediated actin retrograde flow directs neurite formation in the developing braineditor^editors. Neuron. United States: © 2012 Elsevier Inc, p 1091–107.

Furutani Y, Matsuno H, Kawasaki M, Sasaki T, Mori K, Yoshihara Y. 2007. Interaction between telencephalin and ERM family proteins mediates dendritic filopodia formation. J Neurosci 27:8866–76.

Galloway PG, Perry G, Gambetti P. 1987. Hirano body filaments contain actin and actin-associated proteins. J Neuropathol Exp Neurol 46:185–99.

Gardel ML, Sabass B, Ji L, Danuser G, Schwarz US, Waterman CM. 2008. Traction stress in focal adhesions correlates biphasically with actin retrograde flow speed. J Cell Biol 183:999–1005.

Gauthier NC, Masters TA, Sheetz MP. 2012. Mechanical feedback between membrane tension and dynamicseditor^editors. Trends Cell Biol. England: 2012 Elsevier Ltd, p 527–35.

Giasson BI, Mushynski WE. 1997. Developmentally regulated stabilization of neuronal intermediate filaments in rat cerebral cortex. Neurosci Lett 229:77–80.

Gomes C, Cunha C, Nascimento F, Ribeiro JA, Vaz AR, Brites D. 2019. Cortical Neurotoxic Astrocytes with Early ALS Pathology and miR-146a Deficit Replicate Gliosis Markers of Symptomatic SOD1G93A Mouse Modeleditor^editors. Mol Neurobiol. United States, p 2137–2158.

Heo K, Lim SM, Nahm M, Kim YE, Oh KW, Park HT, Ki CS, Kim SH, Lee S. 2018. A De Novo RAPGEF2 Variant Identified in a Sporadic Amyotrophic Lateral Sclerosis Patient Impairs Microtubule Stability and Axonal Mitochondria Distribution. Exp Neurobiol 27:550–563.

Hurtley SM. 1998. Cell biology of the cytoskeleton. Science 279:459.

Ikawa K, Sugimura K. 2018. AIP1 and cofilin ensure a resistance to tissue tension and promote directional cell rearrangement. Nat Commun 9:3295.

Jiu Y, Lehtimäki J, Tojkander S, Cheng F, Jäälinoja H, Liu X, Varjosalo M, Eriksson JE, Lappalainen P. 2015. Bidirectional Interplay between Vimentin Intermediate Filaments and Contractile Actin Stress Fibers. Cell Rep 11:1511–8.

Kwon I, Xiang S, Kato M, Wu L, Theodoropoulos P, Wang T, Kim J, Yun J, Xie Y, McKnight SL. 2014. Poly-dipeptides encoded by the C9orf72 repeats bind nucleoli, impede RNA biogenesis, and kill cells. Science 345:1139–45.

Lagos-Cabré R, Alvarez A, Kong M, Burgos-Bravo F, Cárdenas A, Rojas-Mancilla E, Pérez-Nuñez R, Herrera-Molina R, Rojas F, Schneider P, Herrera-Marschitz M, Quest AFG, van Zundert B, Leyton L 2017. α(V)β(3) Integrin regulates astrocyte reactivity. J Neuroinflammation 14:194.

Leigh PN, Dodson A, Swash M, Brion JP, Anderton BH. 1989. Cytoskeletal abnormalities in motor neuron disease. An immunocytochemical study. Brain 112 (Pt 2):521–35.

Leshchyns’ka I, Sytnyk V. 2016. Synaptic Cell Adhesion Molecules in Alzheimer’s Disease. Neural Plast 2016:6427537.

Lin Y, Mori E, Kato M, Xiang S, Wu L, Kwon I, McKnight SL. 2016. Toxic PR Poly-Dipeptides Encoded by the C9orf72 Repeat Expansion Target LC Domain Polymers. Cell 167:789–802 e12.

Luo L. 2002. Actin cytoskeleton regulation in neuronal morphogenesis and structural plasticityeditor^editors. Annu Rev Cell Dev Biol. United States, p 601–35.

Morris RL, Hollenbeck PJ. 1995. Axonal transport of mitochondria along microtubules and F-actin in living vertebrate neurons. J Cell Biol 131:1315–26.

Nagayama K, Uchida K, Sato A. 2019. A novel micro-grooved collagen substrate for inducing vascular smooth muscle differentiation through cell tissue arrangement and nucleus remodelingeditor^editors. J Mech Behav Biomed Mater. Netherlands: 2018 Elsevier Ltd, p 295–305.

Oakes PW, Beckham Y, Stricker J, Gardel ML. 2012. Tension is required but not sufficient for focal adhesion maturation without a stress fiber template. J Cell Biol 196:363–74.

Osking Z, Ayers JI, Hildebrandt R, Skruber K, Brown H, Ryu D, Eukovich AR, Golde TE, Borchelt DR, Read TA, Vitriol EA. 2019. ALS-Linked SOD1 Mutants Enhance Neurite Outgrowth and Branching in Adult Motor Neurons. iScience 19:448–449.

Parsons JT, Horwitz AR, Schwartz MA. 2010. Cell adhesion: integrating cytoskeletal dynamics and cellular tension. Nat Rev Mol Cell Biol 11:633–43.

Rowland LP, Shneider NA. 2001. Amyotrophic lateral sclerosis. N Engl J Med 344:1688–700.

Sack I, Streitberger KJ, Krefting D, Paul F, Braun J. 2011. The influence of physiological aging and atrophy on brain viscoelastic properties in humans. PLoS One 6:e23451.

Shabbir SH, Cleland MM, Goldman RD, Mrksich M. 2014. Geometric control of vimentin intermediate filaments. Biomaterials 35:1359–66.

Sivadasan R, Hornburg D, Drepper C, Frank N, Jablonka S, Hansel A, Lojewski X, Sterneckert J, Hermann A, Shaw PJ, Ince PG, Mann M, Meissner F, Sendtner M. 2016. C9ORF72 interaction with cofilin modulates actin dynamics in motor neuronseditor^editors. Nat Neurosci. United States, p 1610–1618.

Smith BN, Ticozzi N, Fallini C, Gkazi AS, Topp S, Kenna KP, Scotter EL, Kost J, Keagle P, Miller JW, Calini D, Vance C, Danielson EW, Troakes C, Tiloca C, Al-Sarraj S, Lewis EA, King A, Colombrita C, Pensato V, Castellotti B, de Belleroche J, Baas F, ten Asbroek AL, Sapp PC, McKenna-Yasek D, McLaughlin RL, Polak M, Asress S, Esteban-Perez J, Munoz-Blanco JL, Simpson M, van Rheenen W, Diekstra FP, Lauria G, Duga S, Corti S, Cereda C, Corrado L, Soraru G, Morrison KE, Williams KL, Nicholson GA, Blair IP, Dion PA, Leblond CS, Rouleau GA, Hardiman O, Veldink JH, van den Berg LH, Al-Chalabi A, Pall H, Shaw PJ, Turner MR, Talbot K, Taroni F, Garcia-Redondo A, Wu Z, Glass JD, Gellera C, Ratti A, Brown RH, Jr., Silani V, Shaw CE, Landers JE. 2014. Exome-wide rare variant analysis identifies TUBA4A mutations associated with familial ALS. Neuron 84:324–31.

Webster CP, Smith EF, Bauer CS, Moller A, Hautbergue GM, Ferraiuolo L, Myszczynska MA, Higginbottom A, Walsh MJ, Whitworth AJ, Kaspar BK, Meyer K, Shaw PJ, Grierson AJ, De Vos KJ. 2016. The C9orf72 protein interacts with Rab1a and the ULK1 complex to regulate initiation of autophagy. EMBO J 35:1656–76.

Wu CH, Fallini C, Ticozzi N, Keagle PJ, Sapp PC, Piotrowska K, Lowe P, Koppers M, McKenna-Yasek D, Baron DM, Kost JE, Gonzalez-Perez P, Fox AD, Adams J, Taroni F, Tiloca C, Leclerc AL, Chafe SC, Mangroo D, Moore MJ, Zitzewitz JA, Xu ZS, van den Berg LH, Glass JD, Siciliano G, Cirulli ET, Goldstein DB, Salachas F, Meininger V, Rossoll W, Ratti A, Gellera C, Bosco DA, Bassell GJ, Silani V, Drory VE, Brown RH, Jr., Landers JE. 2012. Mutations in the profilin 1 gene cause familial amyotrophic lateral sclerosis. Nature 488:499–503.

Yabe JT, Chan WK, Wang FS, Pimenta A, Ortiz DD, Shea TB. 2003. Regulation of the transition from vimentin to neurofilaments during neuronal differentiation. Cell Motil Cytoskeleton 56:193–205.

Yamashiro Y, Thang BQ, Ramirez K, Shin SJ, Kohata T, Ohata S, Nguyen TAV, Ohtsuki S, Nagayama K, Yanagisawa H. 2020. Matrix mechanotransduction mediated by thrombospondin-1/integrin/YAP in the vascular remodeling. Proc Natl Acad Sci U S A 117:9896–9905.

Zhang K, Daigle JG, Cunningham KM, Coyne AN, Ruan K, Grima JC, Bowen KE, Wadhwa H, Yang P, Rigo F, Taylor JP, Gitler AD, Rothstein JD, Lloyd TE. 2018. Stress Granule Assembly Disrupts Nucleocytoplasmic Transport. Cell 173:958-971.e17.

Zhang K, Donnelly CJ, Haeusler AR, Grima JC, Machamer JB, Steinwald P, Daley EL, Miller SJ, Cunningham KM, Vidensky S, Gupta S, Thomas MA, Hong I, Chiu SL, Huganir RL, Ostrow LW, Matunis MJ, Wang J, Sattler R, Lloyd TE, Rothstein JD. 2015. The C9orf72 repeat expansion disrupts nucleocytoplasmic transport. Nature 525:56–61.

