## Supplementary figures and images for "*C9orf72*- derived proline:arginine poly-dipeptides disturb cytoskeletal architecture"

### supplementary figure

Supplementary figure 1.

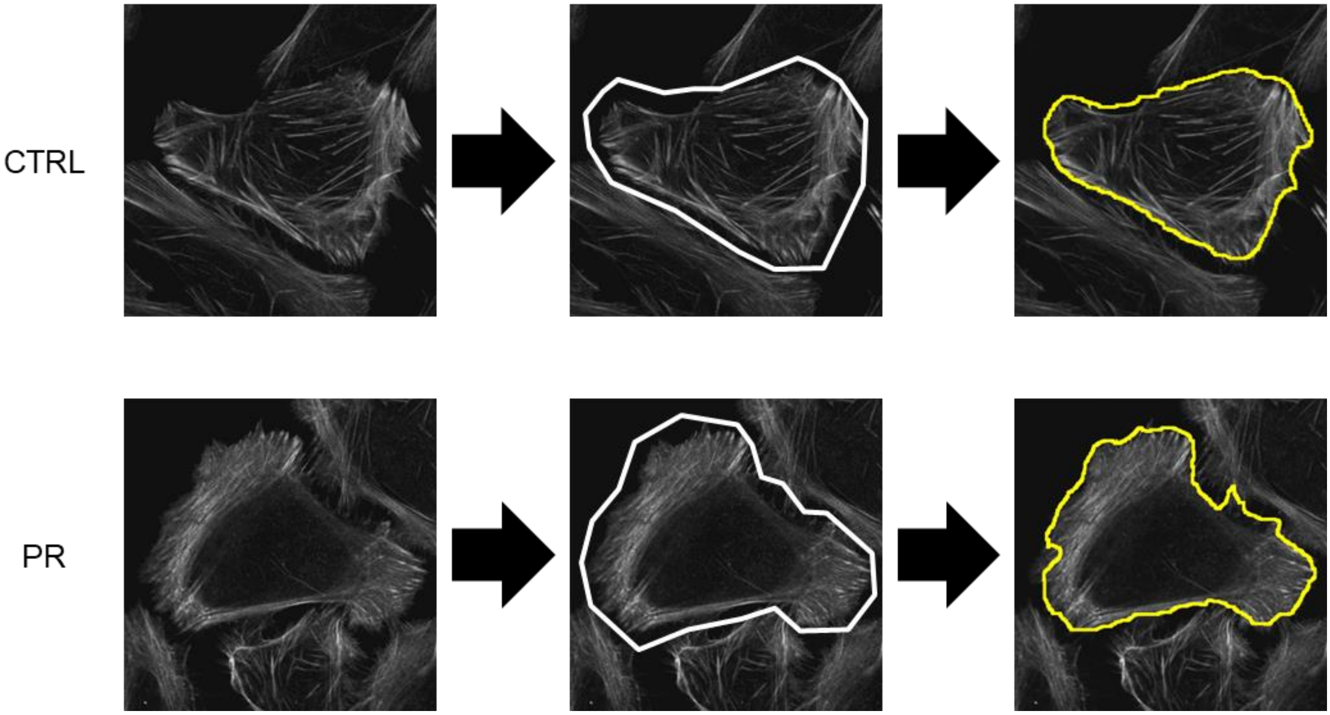

Supplementary figure 2.

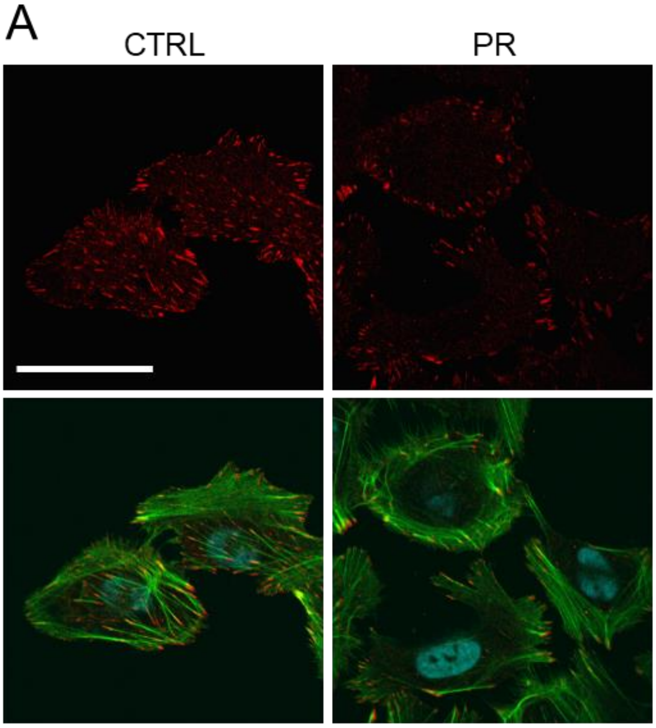


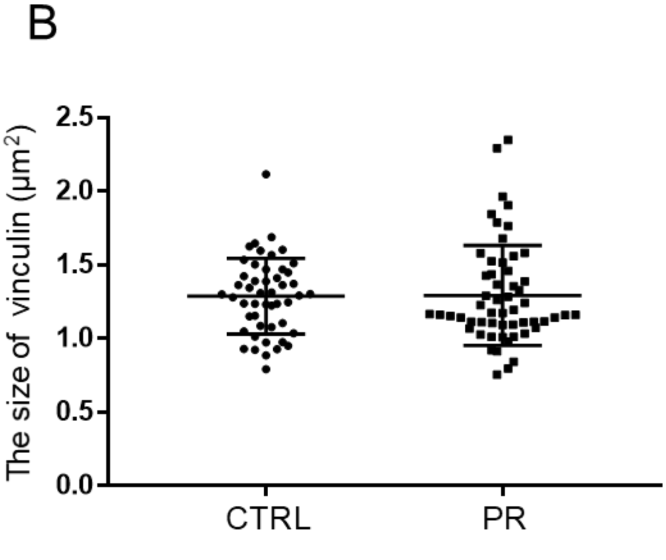
